# A pituitary gene network linking *vgll3* to regulators of sexual maturation in male Atlantic salmon

**DOI:** 10.1101/2022.06.20.496813

**Authors:** Ehsan Pashay Ahi, Marion Sinclair-Waters, Iikki Donner, Craig R. Primmer

**Author notes:** Corresponding Author:* Ehsan Pashay Ahi ^1^.

## Abstract

Age at maturity is a key life history trait and a significant contributor to life history strategy variation. The maturation process is complex and influenced by genetic and environmental factors alike, but specific causes of variation in maturation timing remain elusive. In many species, the increase in the regulatory gonadotropin-releasing hormone 1 (GnRH1) marks the onset of puberty. Atlantic salmon, however, lack the gene encoding GnRH1, suggesting other regulatory factors are involved in the maturation process. Earlier research in Atlantic salmon has found a strong association between alternative alleles of *vgll3* and maturation timing, making *vgll3* a candidate reproductive axis gene regulator. Recently we reported strong induction of gonadotropin encoding genes (*fshb* and *lhb*) in the pituitary of male Atlantic salmon homozygous for the *vgll3* allele linked with the early maturation allele (*E*). The induction of gonadotropins was accompanied by increased expression of their direct upstream regulators, *c-jun* and *sf1* (*nr5a1b*) in the pituitary. In mammals, the transcriptional activation of *c-jun* and *sf1* is also required for induction of *fshb* and *lhb*, however, GnRH1 is responsible for increased transcriptional activity of *c-jun* and *sf1*. The absence of *gnrh1* in salmon raises the possibility of the involvement of other regulators upstream of these factors. In this study, we investigated such a possibility through a stepwise approach for identifying a gene regulatory network (GRN) containing *c-jun* and *sf1* and using the zebrafish coexpression database and transcription factor motif enrichment analysis. We found a GRN containing *c-jun* with predicted upstream regulators, *e2f1, egr1, foxj1* and *klf4*, which are also differentially expressed in the pituitary. Finally, we suggest a model for transcriptional regulation of *c-jun* and *sf1*in the absence of *gnrh1* in the pituitary, which may have broader implications across vertebrates.

## Introduction

Age at maturity, i.e., the age at which an individual reproduces, is a critical trait in an organism’s life history as it affects fitness traits such as survival and reproductive success (Mobley et al., 2021). This life history trait varies both within and among species and thus contributes to the remarkable variation in life history strategies of organisms (Healy et al., 2019). The maturation process is influenced by both genetic and environmental signals and involves changes in, amongst other things, hormone levels and behavior (Varlinskaya et al., 2013), but the specific factors leading to variation in its timing are poorly understood, especially at the molecular level (Howard and Dunkel, 2019; Leka-Emiri et al., 2017; Mobley et al., 2021). The major upstream regulator of sexual maturation in most species is gonadotropin releasing hormone (GnRH), encoded by *gnrh1* and expressed in the hypothalamus (Whitlock et al., 2019). In Atlantic salmon, however, the absence of *gnrh1* in its genome as well as the unresolved compensatory role of other members of the *gnrh* family, *gnrh2* and *gnrh3*, in controlling puberty (Muñoz-Cueto et al., 2020; Whitlock et al., 2019), brings into question the potential role of other molecular regulators.

Genetic variation in *vgll3* (the vestigial-like family member 3 gene) is strongly associated with maturation timing in both sexes of wild Atlantic salmon but also exhibits sex-specific maturation effects (Barson et al., 2015; Czorlich et al., 2018). This association between *vgll3* genotype and maturation probability has been observed in one year-old male parr in common garden settings (Debes et al., 2021; Sinclair-Waters et al., 2021; Verta et al., 2020). An expression study of alternative isoforms of *vgll3* revealed a link between its alleles and variation in age at maturity in one-year-old salmon males (Verta et al., 2020). Our recent study also showed that *vgll3* strongly affects the expression of reproductive axis genes in one year-old males (Ahi et al., 2022). In that study we reported marked increases in transcription of the gonadotropin encoding genes (*fshb* and *lhb*) and two of their upstream transcription factors (TFs), *jun* and *sf1*, in the pituitary of phenotypically immature male Atlantic salmon homozygous for the early-maturing associated vgll3*E allele (Ahi et al., 2022).

In mammals, GnRH1 induces *Fshb* expression by stimulating Ap-1 TF complex, which is formed by a heterodimer of c-Jun (Jun) and c-Fos (Fos) (Coss et al., 2004). GnRH1 particularly enhances the expression of *Jun*, which binds to *Fshb* promoter. Moreover, the expression of *Lhb* and its reproductive function can be dependent on Jun transcriptional activity (Jonak et al., 2018). The second TF, *sf1* (or *nr5a1*), is known to cooperate with *jun* (c-Jun) enhancing transcriptional activity of *jun* in regulating numerous downstream target genes (Dubé et al., 2009; Guo et al., 2007; Martin and Tremblay, 2009). *Sf1* is also a known direct upstream regulator of both *Lhb* and *Fshb* and its transcriptional activity is again under the influence of GnRH1 in mammals (Haisenleder et al., 1996; Kaiser et al., 2000; Keri and Nilson, 1996).

The abovementioned findings in mammals raise the questions how *jun* and *sf1* are induced in the pituitary of Atlantic salmon in the absence of *gnrh1* and also how their transcriptional induction is linked to the vgll3*E allele. Here, we conduct a stepwise approach established in different teleost fish taxa (Ahi et al., 2021, 2015; Ahi and Sefc, 2018; Pohl et al., 2021) that combines knowledge-based and de novo methods with gene expression analysis to identify gene regulatory network(s), or GRNs, linking *vgll3* function to differential expression of *jun* and *sf1* in the absence of *gnrh1* in the pituitary of Atlantic salmon.

## Materials and methods

### Animal material and genotyping

The Atlantic salmon used in this study were a subset of the individuals used in Sinclair-Waters et al. (2021). All individuals used here were males from the same family (FM_T7 in the dryad dataset: https://doi.org/10.5061/dryad.kwh70rz3v). The fish were the offspring of unrelated parents of Kymijoki origin (F1 hatchery generation) that were heterozygous with respect to *vgll3* alleles (*vgll3*EL*) meaning that all three *vgll3* genotypes occurred amongst full sibs of the family. This enabled the assessment of the expression patterns of all the possible *vgll3* genotypes within an otherwise similar genetic background. The *vgll3* genotypes of the offspring were determined as described in Sinclair-Waters et al. (2021).

The fish were euthanized approximately one-year post-fertilization with an overdose of the anesthetic buffered tricaine methane sulfonate (MS-222) and dissected, and sex and maturation status were determined visually by observing the presence of female or male gonads as outlined in Verta et al. (2020). The maturation status of males was classified on a scale from 1 (no phenotypic signs of gonad maturation) to 4 (large gonads leaking milt). All the individuals in the current study were classified as stage 1 and referred to hereafter as “phenotypically immature” (Supplementary data). The gonadosomatic index (GSI) was calculated for a subset of the fish. The GSI for immature category 1 individuals ranged from 0.00 to 0.05, whereas the GSI for mature category 3 and 4 individuals was around 100 times higher, between 4.7-5.6.

### Sample collection, RNA extraction and cDNA synthesis

The brain, pituitary, and testes of 34 (ten *vgll3*EE*, 16 *vgll3*EL*, and eight *vgll3*LL*) phenotypically immature males were sampled immediately following euthanasia using fine forceps. Tissue samples were snap frozen in liquid nitrogen and stored at -80 °C until extraction. Dissected tissues were homogenized with a bead mill homogenizer, Bead Ruptor Elite (Omni International Inc.). We extracted RNA from the sampled brain and testis using the NucleoSpin RNA kit (Macherey-Nagel GmbH & Co. KG), and the pituitary using the NucleoSpin RNA Clean-up XS kit (Macherey-Nagel GmbH & Co. KG). RNA samples were eluted in 50 μl nuclease-free water, and their concentration and quality assessed with NanoDrop ND-1000 and the 2100 Bioanalyzer system (Agilent Technologies, Inc.). To synthesize cDNA, we used 500 ng of RNA per sample and the iScript cDNA Synthesis Kit (Bio-Rad Laboratories, Inc.).

### Gene selection, designing primers and qPCR

We followed a previously established approach for GRN identification (Ahi et al., 2021, 2015; Ahi and Sefc, 2018; Pohl et al., 2021) using zebrafish co-expression data available at COXPRESdb (http://coxpresdb.jp/) version 7.0 (Obayashi et al., 2019). To do this, we first selected the 10 genes with the highest co-expression values with *jun* and *sf1* (*nr5a1b*). Among the tested candidate co-expressed genes, those exhibiting expression differences similar to *jun* were selected for the upstream regulator prediction step. We conducted motif enrichment on 4 kb upstream sequences (promoter and 5’-UTR) of these genes using the MEME Suite algorithm (Bailey et al., 2009). The overrepresented motifs in the upstream regulatory sequences of the genes were compared to position weight matrices (PWMs) attained from the TRANSFAC database (Matys et al., 2003) using the STAMP tool (Mahony and Benos, 2007) in order to predict potential transcription factor (TF) binding sites. The prediction of functional associations was conducted using STRING v10 (http://string-db.org/), an interactome databases for vertebrates.

Primer design was as described in Ahi et al., 2019, using two online tools: Primer Express 3.0 (Applied Biosystems, CA, USA) and OligoAnalyzer 3.1 (Integrated DNA Technology) (Supplementary data) and gene sequences obtained from the recently annotated *Salmo salar* genome in the Ensembl database, http://www.ensembl.org. The qPCR reactions were prepared as described in Ahi et al., 2019, using PowerUp SYBR Green Master Mix (Thermo Fischer Scientific), and performed on a Bio-Rad CFX96 Touch Real Time PCR Detection system (Bio-Rad, Hercules, CA, USA). The details of the qPCR program and calculation of primer efficiencies are described in Ahi et al., 2019.

### Analysis of gene expression data

To calculate the expression levels of our target genes, we utilized three reference genes validated in the pituitary of Atlantic salmon: *hprt1, gapdh*, and *elf1a*. The geometric mean of the Cq values of of these genes was used as the normalization factor together with the following formula: ΔCq _target_ = Cq _target_ – Cq _reference_. For each gene, a biological replicate with the lowest expression level across all the samples of the three *vgll3* genotypes (calibrator sample) was selected to calculate ΔΔCq values (ΔCq _target_ – ΔCq _calibrator_). The relative expression quantities (RQ values) were calculated as 2^−ΔΔCq^, and their fold changes (logarithmic values of RQs) were used for statistical analysis (Pfaffl, 2001). The Student’s t-test was applied for direct comparisons of gene expression levels between the genotypes, followed by Benjamini-Hochberg correction for multiple comparisons (Thissen et al., 2002).

## Results

### Expression patterns of *nr5a1b* co-expressed candidate genes

Expression analysis of the top 10 genes co-expressed with *nr5a1b* revealed that only 3 genes, *efna5b, mtf1* and *zgc:113142* had marginally significant expression level differences between genotypes (Figure 1). These differences were unlike our previous finding in which *nr5a1b* expression was much higher in the vgll3*EE genotype than EL and LL genotypes. More specifically, *enfa5b* only shows a significant difference between EE and EL genotypes and *mtf1* displays a difference between the EE and LL genotypes, whereas *zgc:113142* shows opposite expression patterns to *nr5a1b*, i.e. LL and EL genotypes have higher expression than the EE genotype. These observations suggest the absence of a co-regulatory connection between *nr5a1b* and the selected co-expressed genes in the pituitary of Atlantic salmon. This may also indicate that expression differences in *nr5a1b* are controlled by a regulatory element unique to this gene, which is probably not present in the promoters of the other co-expressed genes.

**Figure 1.**
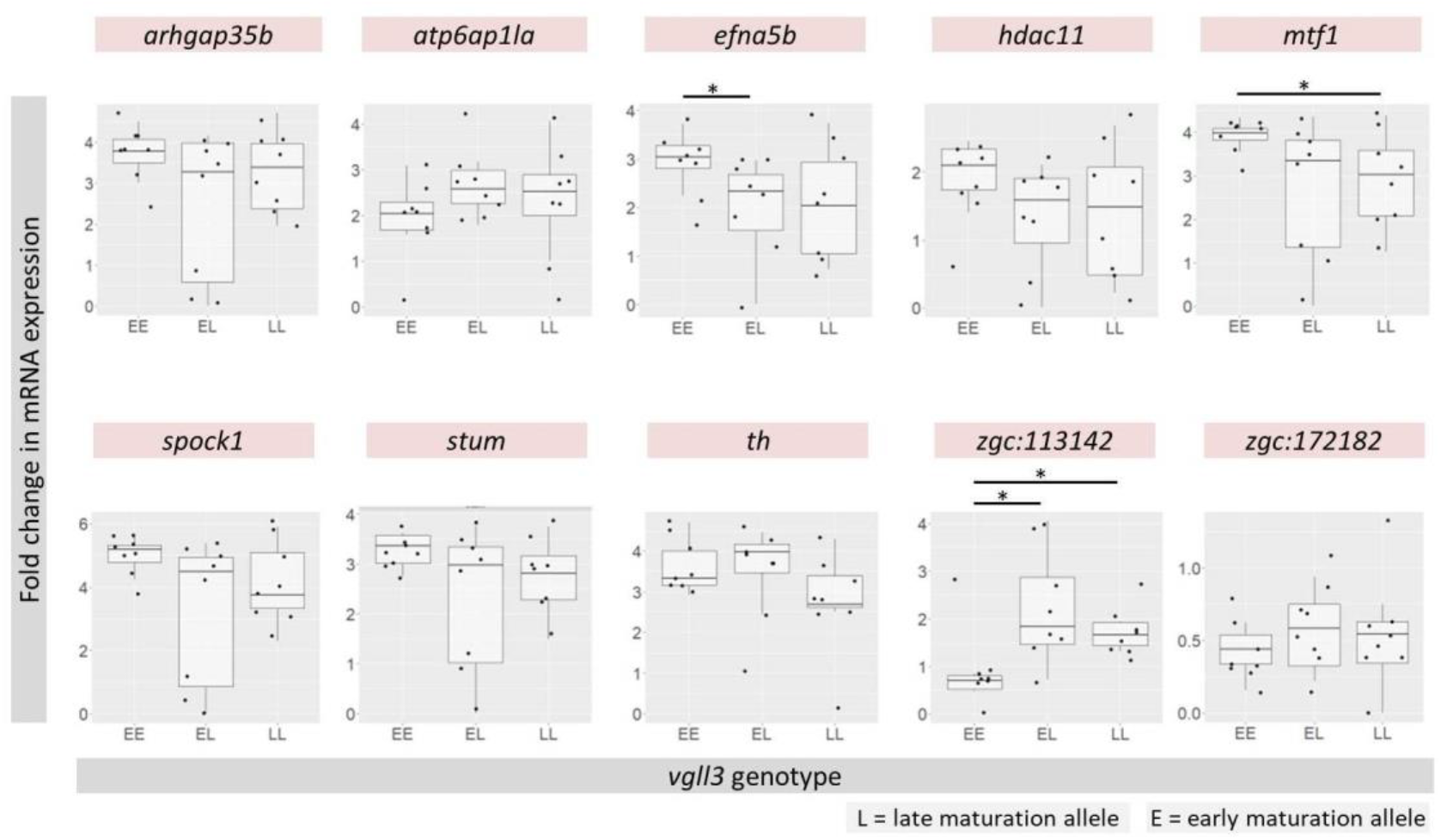
Expression analysis of selected candidate genes co-expressed with *nr5a1b* in the pituitary of male Atlantic salmon with different *vgll3* genotypes. Small dots indicate individual expression levels, the middle line represents the median, the box indicates the 25/75 percentiles and the whiskers the 5/95 percentiles for each plot. Asterisks above each plot indicate significantly different differential expression comparisons (* P < 0.05; ** P < 0.01; *** P < 0.001).

### Expression patterns of *jun* co-expressed candidate genes

We implemented the same approach to identify differential expression in *jun* co-expressed genes by selecting the top 10 genes ranked with highest probability of expression correlation with *jun*. Strikingly, all the selected genes were found to display significantly higher expression in vgll3*EE genotype individuals than in individuals with the two other *vgll3* genotypes (Figure 2), thus following a similar expression pattern observed for *jun* in our previous study (Ahi et al., 2022). These findings provide evidence supporting the possible presence of a co-regulatory connection between *jun* and the selected co-expressed genes in the pituitary of Atlantic salmon.

**Figure 2.**
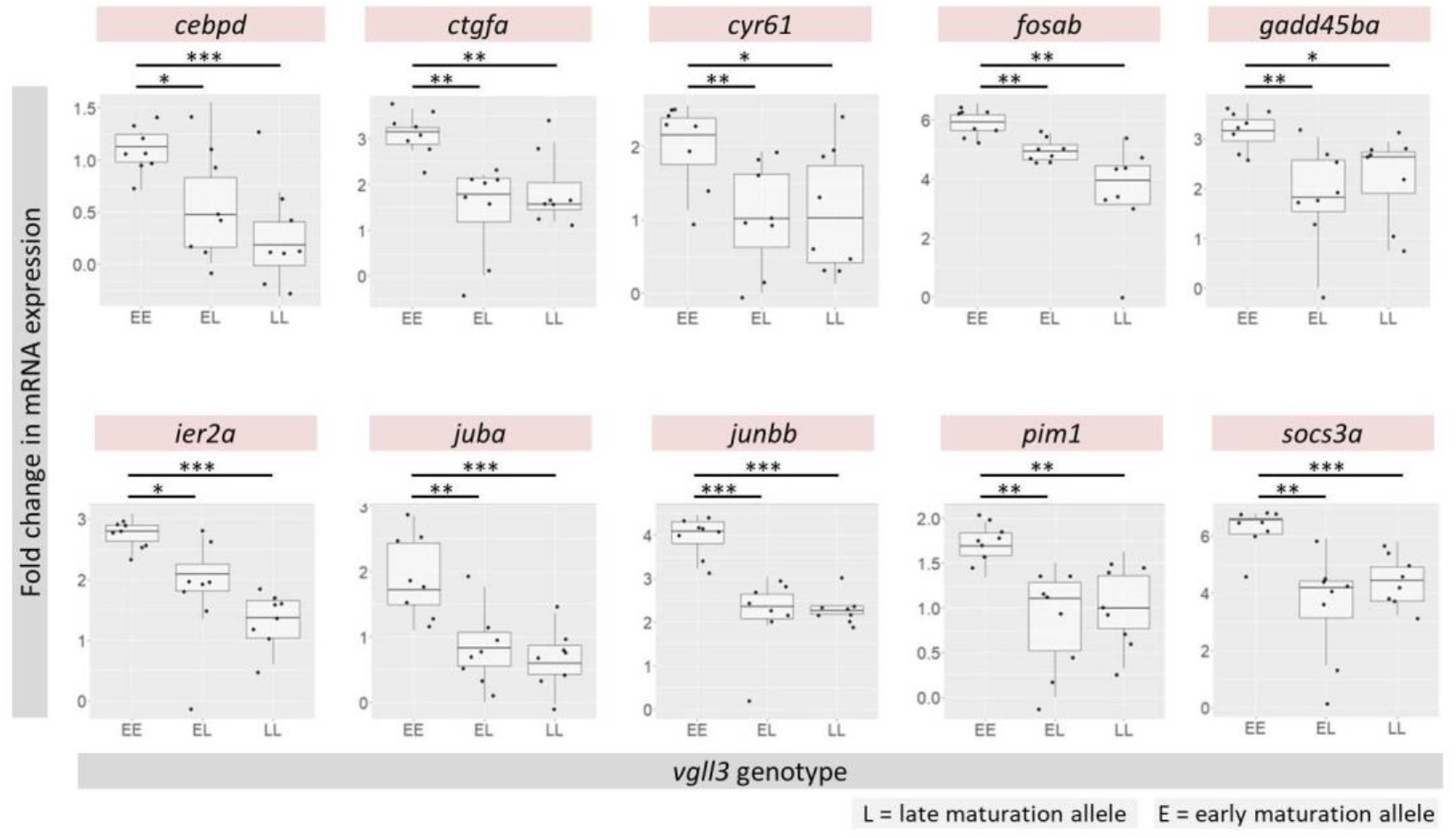
Expression analysis of selected candidate genes co-expressed with *jun* in the pituitary of male Atlantic salmon with different *vgll3* genotypes. Small dots indicate individual expression levels, the middle line represents the median, the box indicates the 25/75 percentiles and the whiskers the 5/95 percentiles for each plot. Asterisks above each plot indicate the level of significance for differential expression comparisons (* P < 0.05; ** P < 0.01; *** P < 0.001).

### Identification of transcription factors at upstream of *jun* co-expression module

Next, we explored the presence of potential upstream regulators of the identified *jun* co-expression module by investigating *de novo* enrichment of TF binding sites in the upstream regulatory sequences of the co-expressed genes. To do this, we used the promoters and 5’-UTR sequences of the *jun* co-expression module, i.e. *jun* and the top 10 candidate co-expressed genes with a similar expression patterns to *jun* (see the above section), for the motif enrichment analysis step. We identified 8 motifs enriched in the regulatory sequences of almost all of these genes (Table 1). We then parsed the motifs against the database for vertebrate TF binding sites and listed the top matched TF(s) for each motif in Table 1.

**Table 1.**
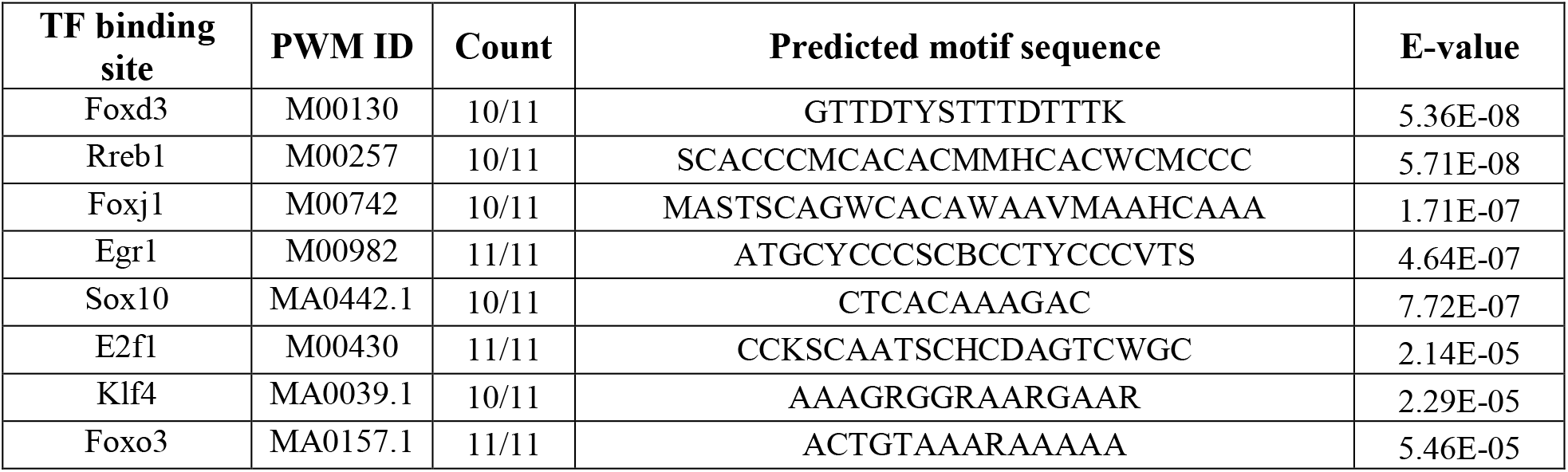
Predicted TF binding sites for potential upstream regulators of the identified *jun* co-expressed genes. PWD ID indicates the positional weight matrix ID of a predicted binding site and E-values refer to matching similarity between the predicted motif sequences and the PWD IDs. ‘Count’ indicates the number of genes inferred in the network containing the predicted motif sequence in their regulatory region.

### Expression patterns of predicted TFs at upstream of of *jun* co-expression module

The expression analysis of potential upstream TFs of the *jun* co-expression module showed that four of the TFs, *e2f1, egr1, klf4* and *foxj1a*, had similar expression patterns to *jun* co-expression module genes, i.e. higher expression in *vgll3\**EE individuals than *vgll3\**EL and *vgll3\**LL individuals (Figure 3). It should be noted that the expression of one of the predicted TFs, *foxd3*, was not detected by qPCR, possibly due to its very low expression level in the pituitary. Taken together, these findings suggest that the GRN consists of four TFs upstream of a co-expression module of genes containing *jun* itself, and that the transcriptional activity of the GRN is under the influence of different *vgll3* alleles. Finally in order to find functional links between *vgll3* and the four TFs, we conducted functional associations analyses of the TFs together with *vgll3* and other components of the Hippo pathway (yap1, TAZ and TEADs) using an interactome database for vertebrates (Szklarczyk et al., 2015) (Figure 4A). The result showed that the TFs only have regulatory connections with yap1 and TEADs but not vgll3.

**Figure 3.**
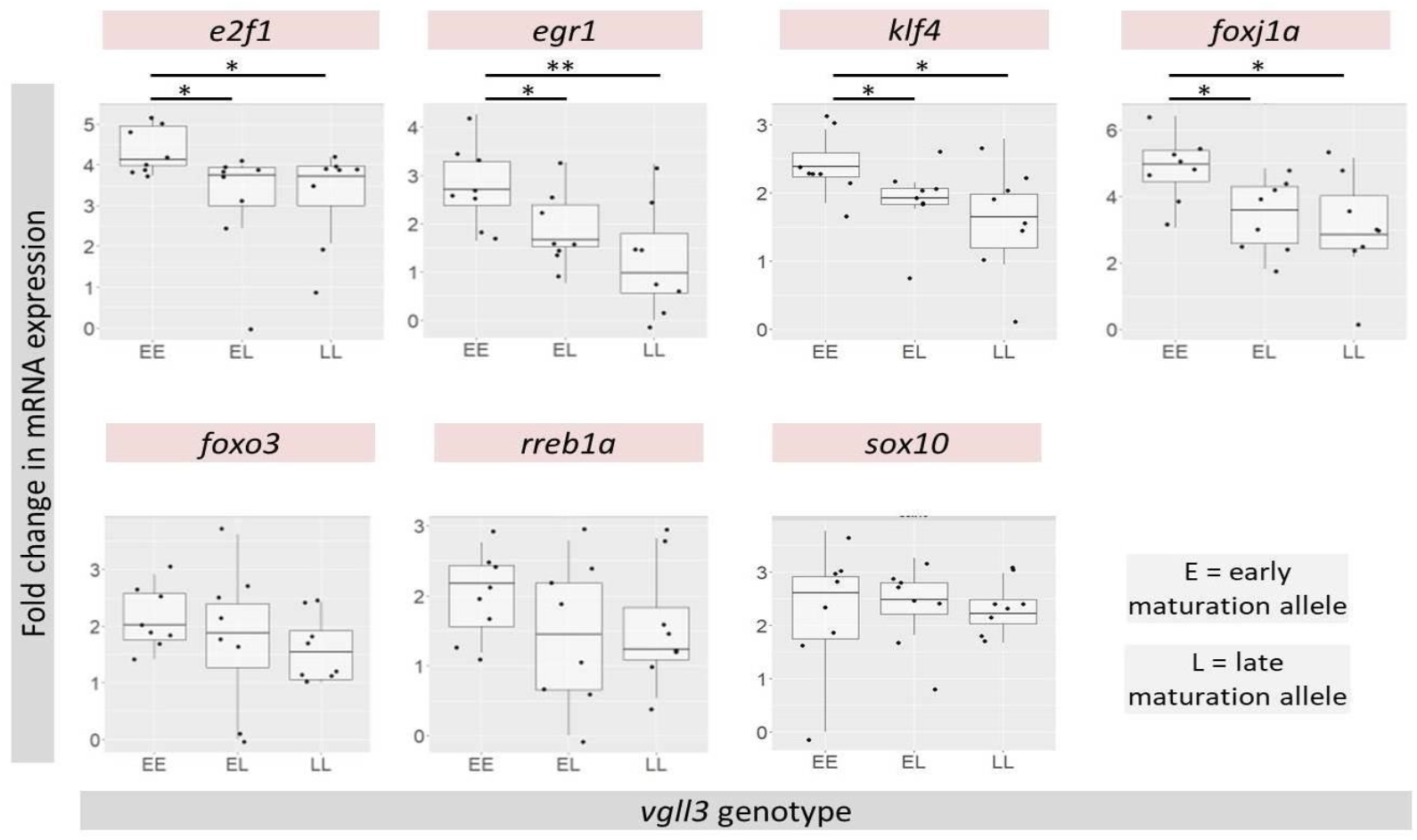
Expression analysis of predicted upstream regulators of *jun* co-expressed genes in the pituitary of male Atlantic salmon with different *vgll3* genotypes. Small dots indicate individual expression levels, the middle line represents the median, the box indicates the 25/75 percentiles and the whiskers the 5/95 percentiles for each plot. Asterisks above each plot indicate the level of significance for differential expression comparison (* P < 0.05; ** P < 0.01; *** P < 0.001).

**Figure 4.**
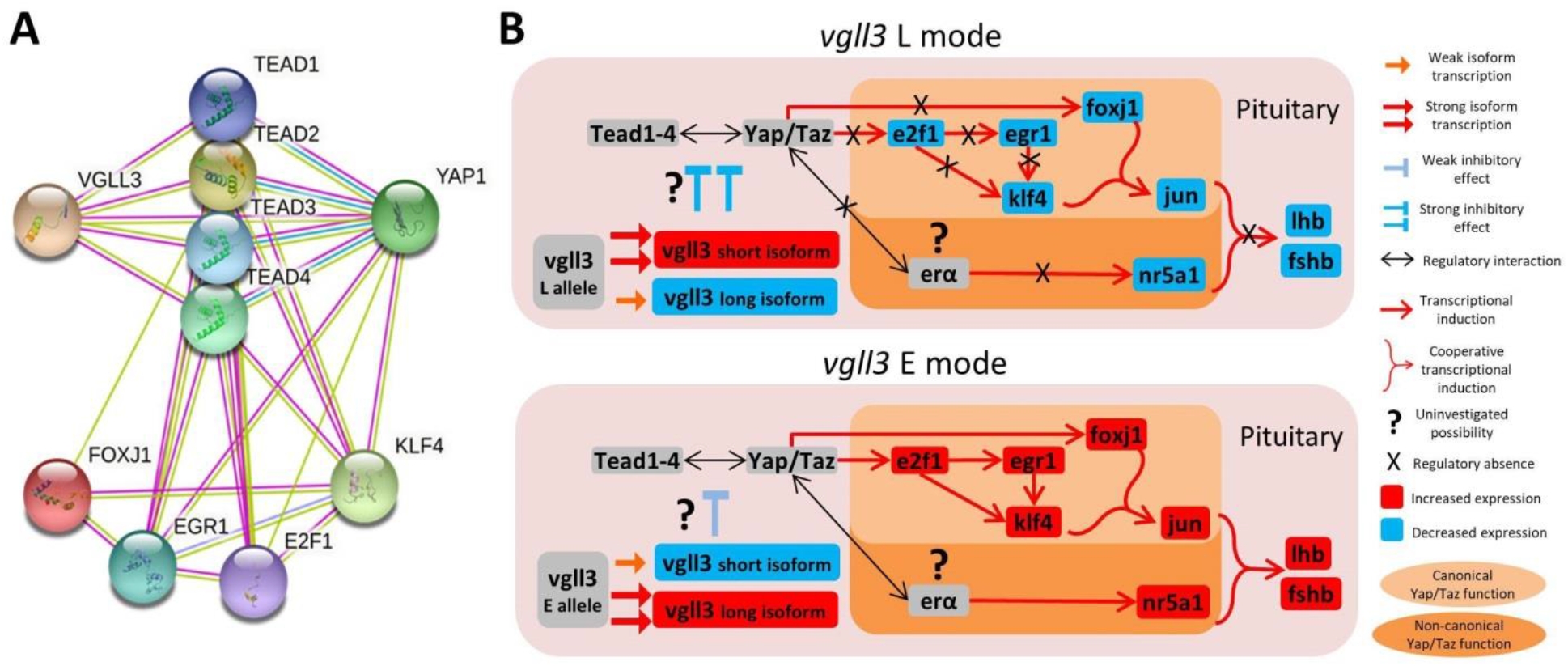
Alternative modes of *vgll3*-allele regulatory effects on upstream regulators of gonadotropins in male Atlantic salmon. (**A**) Predicted functional associations between the predicted differentially expressed TFs and components of the Hippo signaling pathway based on STRING v10 (http://string-db.org/), an interactome databases for vertebrates. (**B**) Proposed models for the indirect *vgll3* regulatory connections with the genes investigated in this study. We hypothesize that the *vgll3*\*L allele, which promotes transcription of *vgll3* short isoform with stronger inhibitory effects on Yap/Taz binding to Tead1-4, is responsible for the decreased expression level of the identified transcription factors, *e2f1, egr1, foxj1* and *klf4*. In the *vgll3*\*E allele, however, the short isoform of vgll3 is less transcribed causing more Yap/Taz binding to Tead1-4. This results transcriptional activation of Yap/Taz downstream targets i.e. the identified TFs, and thus also *jun* and gonadotropin encoding genes.

## Discussion

In this study, we combined knowledge-based and *de novo* methods with gene expression analysis to identify GRNs, linking *vgll3* function to differential expression of *jun* and *sf1* in the absence of *gnrh1* in the pituitary of Atlantic salmon. Using this approach, all 10 of the *jun* co-expressed candidate genes assessed displayed significantly higher expression in *vgll3*EE* genotype individuals than in individuals with the two other *vgll3* genotypes. This *vgll3* genotype associated expression pattern is in a similar direction to that observed earlier for *jun* (Ahi et al., 2022) indicating the potential existence of shared upstream factors regulating the gene network. In addition we predicted 4 TFs, *e2f1, egr1, foxj1* and *klf4*, upstream of the co-expressed genes that showed a similar expression pattern to *jun* with higher expression in vgll3*EE genotypes. Such a co-regulatory connection could be the result of transcriptional activity of shared upstream regulators binding to similar sites on the promoters or short distance sequences of the *jun* co-expressed genes.

Previous research in mammals provides some clues as to how these four transcription factors could potentially interact and subsequently exert their transcriptional activity in a *vgll3* genotype-specific manner. For example, *e2f1* expression induction might be responsible for the increased expression of *klf4* and *egr1* (Riverso et al., 2017; Zhang et al., 2014; Zheng et al., 2009), but not *foxj1* in the pituitary of the *vgll3*EE* genotype. Moreover, it has been shown that *egr1* can act directly upstream of *klf4* and induces its expression (Lai et al., 2012; Riddick et al., 2017). On the other hand, the similar expression patterns of the four identified TFs here could also indicate cooperative interactions among them in transcriptional regulation of *jun* and the other downstream co-expressed genes. In mammalian cells, cooperative interactions in regulation of shared downstream target genes have already been reported between *egr1* and *klf4* (Whitlock et al., 2011), and between *e2f1* and *egr1* (Wu et al., 2017).

Direct connections between *vgll3* and the abovementioned TFs have not been reported in any vertebrates, however, an indirect connection is between these factors and *vgll3* is conceivable through components of the Hippo signaling pathway. Based on findings in Atlantic salmon and other vertebrates, a multi-level connection between *vgll3* and other components of the Hippo pathway has been recently proposed (Kurko et al., 2020). Vgll3 binds to TEAD transcription factors, which are major regulators of Hippo signaling, and acts as a cofactor influencing the downstream effects of the Hippo signaling pathway (Figeac et al., 2019; Hori et al., 2020; Kjærner-Semb et al., 2018; Meng et al., 2016; Mesrouze et al., 2020). More specifically, vgll3 can activate the Hippo pathway by competing with YAP/TAZ interaction with TEADs, thereby, suppressing genes downstream of YAP/TAZ/TEADs while enhancing transcription of vgll3/TEADs downstream target genes (Hori et al., 2020). In Atlantic salmon, it has been shown that expression of *vgll3* is negatively correlated with *yap*, which indicates negative transcriptional regulatory effects between these cofactors (Kurko et al., 2020). In addition, a recent study hypothesized that a short isoform of *vgll3*, mainly encoded by the *vgll3*L* allele, has stronger inhibitory effects on *yap1* expression and its interaction with TEADs (Verta et al., 2020). By predicting functional associations of the identified TFs, along with vgll3, yap1 and TEADs, using an interactome database for vertebrates (Szklarczyk et al., 2015), our current study found that the TFs only show regulatory connections with yap1 and TEADs but not vgll3 (Figure 4A). This could indicate an indirect regulatory connection between vgll3 and the identified TFs through yap1 and TEADs. Interestingly, it has been shown in mammalian cells that Yap acts directly upstream of two of the identified TFs, *e2f1* and *foxj1*, by enhancing their transcription (Mahoney et al., 2014; Mizuno et al., 2012; Oku et al., 2018). These findings in mammals could explain both an indirect connection between the identified TFs and vgll3, as well as the reduced expression of the TFs in the pituitary of fish carrying the *vgll3*L* allele. Using this knowledge, we propose a hierarchical regulatory cascade in which the short *vgll3* isoform has stronger inhibitory effects on *yap1* and thus leads to reduced expression of *yap1* and its downstream TFs, *e2f1* and *foxj1*. Subsequently, the decreased *e2f1* expression leads to reduced expression of its downstream target TFs, *egr1* and *klf4*. Finally, the reduced expression of all four TFs predicted upstream of *jun* causes reduced expression of *jun* and its co-expressed genes (depicted in Figure 4B).

By combining knowledge of previous studies with our current results, we propose another hierarchical regulatory cascade involving Hippo signaling, *sf1*, and gonadotropins in the pituitary of male Atlantic salmon, which may be affected by changes in the availability of different *vgll3* isoforms (Fig. 4B). In this cascade, the stronger vgll3 mediated repression of the Yap/Taz interaction with Tead1-4 in the *vgll3*LL* genotype weakens the stimulatory effect of Yap/Taz/Tead on a potential ERα-activated enhancer (at upstream of *sf1*). Subsequently, this leads to reduced expression of *sf1* in the pituitary of *vgll3*LL* compared to *vgll3*EE* individuals (see Figure 4). *Sf1* (also called *nr5a1b*) is a TF belonging to the nuclear receptor superfamily, and is essential for anterior pituitary stem cell differentiation into the gonadotrope cell lineage (Ingraham et al., 1994). As with *jun*, we tested the expression of the top genes co-expressed with *sf1* but did not find similar expression differences. This could indicate that regulatory elements other than those acting on short distance binding elements (within the promoters) may be involved, since the co-expressed genes usually have binding sites for similar TFs on their promoter region. One possibility could be the existence of a long distance enhancer specifically regulating *sf1* expression (but not the other co-expressed genes). Evidence in mammals supports this hypothesis: a pituitary ERα-activated enhancer (an enhancer for the estrogen receptor) has been shown to specifically trigger the expression of *Sf1* in mammals (Pacini et al., 2019). It has been shown in mammalian cells that ERα enhancers can be activated through a non-canonical YAP/TEAD pathway by which both YAP and TEAD act as co-factors binding to ERα enhancers and increase estrogen-induced transcriptional activity (Zhu et al., 2019).

Taken together, we propose a model which summarizes our results and hypothesis by depicting potential hierarchical and interconnected GRN involving Hippo signaling, *jun* and *sf1* TFs and gonadotropins in the pituitary of male Atlantic salmon, which may be affected by changes in the changes availability of different *vgll3* isoforms (Fig. 4B). Moreover, the proposed regulatory connections seem to be independent of *gnrh1* function, which is lacking in the genome Atlantic salmon. Future studies with larger sample sizes including good representation of all *vgll3* genotypes as well as different population backgrounds will help shed further light on the details and expression patterns of the proposed GRN.

## Conclusions

This study provides the first evidence for the existence of a complex gene regulatory network in the pituitary of Atlantic salmon that appears to be downstream of the Hippo pathway, i.e. under direct control of Yap/TEADs canonical and non-canonical signals, and thus, indirectly linked to *vgll3* function. The canonical signal consists of transcription factors such as *e2f1, egr1, klf4* and *foxj1a* and their downstream genes, *jun* and its co-expressed genes, whereas the non-canonical signal involves *sf1* (*nr5a1b*) probably through an ERα-activated enhancer. Both signals seem to be active in transcriptional regulation of gonadotropin encoding genes. The gene regulatory network proposed here may be one of the molecular cascades that regulate transcription of gonadotropins in species lacking *gnrh1*. Further functional investigations are required to validate this hypothesis.

## Acknowledgements

We acknowledge Jaakko Erkinaro and staff at the Natural Resources Institute Finland (Luke) hatchery in Laukaa and members of the Evolution, Conservation and Genomics research group for their help in coordinating and collecting gametes for crosses. We thank Jukka-Pekka Verta for valuable discussions and Nikolai Piavchenko and a number of other assistants for help with fish husbandry, Jacqueline Moustakas-Verho and Shadi Janouz for dissection assistance as well as Morgane Frapin, Seija Tillanen and Annukka Ruokolainen for laboratory assistance.

## Author Contributions

EP, CRP, MS-W conceived the study; EP, ID, MS-W performed experiments; EP developed methodology and analyzed the data; EP, CRP, ID, MS-W interpreted results of the experiments; EP, CRP, ID drafted the manuscript, with EP having the main contribution, and all authors approved the final version of manuscript.

## Funding Source Declaration

Funding was provided by Academy of Finland (grant numbers 307593, 302873, 327255 and 342851), the European Research Council under the European Articles Union’s Horizon 2020 research and innovation program (grant no. 742312) and a Natural Sciences and Engineering Research Council of Canada postgraduate scholarship.

## Competing financial interests

Authors declare no competing interests

## Ethical approval

Animal experimentation followed European Union Directive 2010/63/EU under license ESAVI/42575/2019 granted by the Animal Experiment Board in Finland (ELLA).

## Data availability

All the gene expression data generated during this study are included in this article.

## Notes

### Competing Interest Statement

The authors have declared no competing interest.

## References

Ahi, E.P., Richter, F., Lecaudey, L.A., Sefc, K.M., 2019. Gene expression profiling suggests differences in molecular mechanisms of fin elongation between cichlid species. Sci. Rep. 9. https://doi.org/10.1038/s41598-019-45599-w

Ahi, E.P., Sefc, K.M., 2018. Towards a gene regulatory network shaping the fins of the Princess cichlid. Sci. Rep. 8, 9602. https://doi.org/10.1038/s41598-018-27977-y

Ahi, E.P., Sinclair-Waters, M., Moustakas-Verho, J., Jansouz, S., Primmer, C.R., 2022. Strong regulatory effects of vgll3 genotype on reproductive axis gene expression in juvenile male Atlantic salmon. Gen. Comp. Endocrinol. 325, 114055. https://doi.org/10.1016/j.ygcen.2022.114055

Ahi, E.P., Steinhäuser, S.S., Pálsson, A., Franzdóttir, S.R., Snorrason, S.S., Maier, V.H., Jónsson, Z.O., 2015. Differential expression of the aryl hydrocarbon receptor pathway associates with craniofacial polymorphism in sympatric Arctic charr. Evodevo 6. https://doi.org/10.1186/s13227-015-0022-6

Ahi, E.P., Tsakoumis, E., Brunel, M., Schmitz, M., 2021. Transcriptional study reveals a potential leptin-dependent gene regulatory network in zebrafish brain. Fish Physiol. Biochem. https://doi.org/10.1007/s10695-021-00967-0

Ayllon, F., Solberg, M.F., Glover, K.A., Mohammadi, F., Kjærner-Semb, E., Fjelldal, P.G., Andersson, E., Hansen, T., Edvardsen, R.B., Wargelius, A., 2019. The influence of vgll3 genotypes on sea age at maturity is altered in farmed mowi strain Atlantic salmon. BMC Genet. 20, 44. https://doi.org/10.1186/s12863-019-0745-9

Bailey, T.L., Boden, M., Buske, F.A., Frith, M., Grant, C.E., Clementi, L., Ren, J., Li, W.W., Noble, W.S., 2009. MEME SUITE: tools for motif discovery and searching. Nucleic Acids Res. 37, W202–8. https://doi.org/10.1093/nar/gkp335

Barson, N.J., Aykanat, T., Hindar, K., Baranski, M., Bolstad, G.H., Fiske, P., Jacq, C., Jensen, A.J., Johnston, S.E., Karlsson, S., Kent, M., Moen, T., Niemelä, E., Nome, T., Næsje, T.F., Orell, P., Romakkaniemi, A., Sægrov, H., Urdal, K., Erkinaro, J., Lien, S., Primmer, C.R., 2015. Sex-dependent dominance at a single locus maintains variation in age at maturity in salmon. Nature 528, 405–408. https://doi.org/10.1038/nature16062

Coss, D., Jacobs, S.B.R., Bender, C.E., Mellon, P.L., 2004. A Novel AP-1 Site Is Critical for Maximal Induction of the Follicle-stimulating Hormone β Gene by Gonadotropin-releasing Hormone. J. Biol. Chem. 279, 152–162. https://doi.org/10.1074/jbc.M304697200

Czorlich, Y., Aykanat, T., Erkinaro, J., Orell, P., Primmer, C.R., 2018. Rapid sex-specific evolution of age at maturity is shaped by genetic architecture in Atlantic salmon. Nat. Ecol. Evol. 2, 1800–1807. https://doi.org/10.1038/s41559-018-0681-5

Debes, P. V, Piavchenko, N., Ruokolainen, A., Ovaskainen, O., Moustakas-Verho, J.E., Parre, N., Aykanat, T., Erkinaro, J., Primmer, C.R., 2021. Polygenic and major-locus contributions to sexual maturation timing in Atlantic salmon. Mol. Ecol. https://doi.org/10.1111/MEC.16062

Dubé, C., Bergeron, F., Vaillant, M.J., Robert, N.M., Brousseau, C., Tremblay, J.J., 2009. The nuclear receptors SF1 and LRH1 are expressed in endometrial cancer cells and regulate steroidogenic gene transcription by cooperating with AP-1 factors. Cancer Lett. 275, 127–138. https://doi.org/10.1016/j.canlet.2008.10.008

Figeac, N., Mohamed, A.D., Sun, C., Schönfelder, M., Matallanas, D., Garcia-Munoz, A., Missiaglia, E., Collie-Duguid, E., De Mello, V., Pobbati, A. V., Pruller, J., Jaka, O., Harridge, S.D.R., Hong, W., Shipley, J., Vargesson, N., Zammit, P.S., Wackerhage, H., 2019. VGLL3 operates via TEAD1, TEAD3 and TEAD4 to influence myogenesis in skeletal muscle. J. Cell Sci. 132. https://doi.org/10.1242/jcs.225946

Guo, I.-C., Huang, C.-Y., Wang, C.-K.L., Chung, B., 2007. Activating Protein-1 Cooperates with Steroidogenic Factor-1 to Regulate 3′,5′-Cyclic Adenosine 5′-Monophosphate-Dependent Human CYP11A1 Transcription in Vitro and in Vivo. Endocrinology 148, 1804–1812. https://doi.org/10.1210/en.2006-0938

Haisenleder, D.J., Yasin, M., Dalkin, A.C., Gilrain, J., Marshall, J.C., 1996. GnRH Regulates Steroidogenic Factor-1 (SF-I) Gene Expression in the Rat Pituitary. Endocrinology 137, 5719–5722. https://doi.org/10.1210/endo.137.12.8940405

Healy, K., Ezard, T.H.G., Jones, O.R., Salguero-Gómez, R., Buckley, Y.M., 2019. Animal life history is shaped by the pace of life and the distribution of age-specific mortality and reproduction. Nat. Ecol. Evol. 3, 1217–1224. https://doi.org/10.1038/s41559-019-0938-7

Hori, N., Okada, K., Takakura, Y., Takano, H., Yamaguchi, Naoto, Yamaguchi, Noritaka, 2020. Vestigial-like family member 3 (VGLL3), a cofactor for TEAD transcription factors, promotes cancer cell proliferation by activating the Hippo pathway. J. Biol. Chem. 295, 8798–8807. https://doi.org/10.1074/jbc.ra120.012781

Howard, S.R., Dunkel, L., 2019. Delayed Puberty - Phenotypic Diversity, Molecular Genetic Mechanisms, and Recent Discoveries. Endocr. Rev. https://doi.org/10.1210/er.2018-00248

Ingraham, H.A., Lala, D.S., Ikeda, Y., Luo, X., Shen, W.H., Nachtigal, M.W., Abbud, R., Nilson, J.H., Parker, K.L., 1994. The nuclear receptor steroidogenic factor 1 acts at multiple levels of the reproductive axis. Genes Dev. 8, 2302–2312. https://doi.org/10.1101/gad.8.19.2302

Jonak, C.R., Lainez, N.M., Boehm, U., Coss, D., 2018. GnRH Receptor Expression and Reproductive Function Depend on JUN in GnRH Receptor−Expressing Cells. Endocrinology 159, 1496–1510. https://doi.org/10.1210/en.2017-00844

Kaiser, U.B., Halvorson, L.M., Chen, M.T., 2000. Sp1, Steroidogenic Factor 1 (SF-1), and Early Growth Response Protein 1 (Egr-1) Binding Sites Form a Tripartite Gonadotropin-Releasing Hormone Response Element in the Rat Luteinizing Hormone-β Gene Promoter: an Integral Role for SF-1. Mol. Endocrinol. 14, 1235–1245. https://doi.org/10.1210/mend.14.8.0507

Keri, R.A., Nilson, J.H., 1996. A steroidogenic factor-1 binding site is required for activity of the luteinizing hormone β subunit promoter in gonadotropes of transgenic mice. J. Biol. Chem. 271, 10782–10785. https://doi.org/10.1074/jbc.271.18.10782

Kjærner-Semb, E., Ayllon, F., Kleppe, L., Sørhus, E., Skaftnesmo, K., Furmanek, T., Segafredo, F.T., Thorsen, A., Fjelldal, P.G., Hansen, T., Taranger, G.L., Andersson, E., Schulz, R.W., Wargelius, A., Edvardsen, R.B., 2018. Vgll3 and the Hippo pathway are regulated in Sertoli cells upon entry and during puberty in Atlantic salmon testis. Sci. Rep. 8, 1–11. https://doi.org/10.1038/s41598-018-20308-1

Kurko, J., Debes, P. V, House, A.H., Aykanat, T., Erkinaro, J., Primmer, C.R., 2020. Transcription Profiles of Age-at-Maturity-Associated Genes Suggest Cell Fate Commitment Regulation as a Key Factor in the Atlantic Salmon Maturation Process. G3 Genes|Genomes|Genetics 10, 235–246. https://doi.org/10.1534/g3.119.400882

Lai, J.K., Wu, H.C., Shen, Y.C., Hsieh, H.Y., Yang, S.Y., Chang, C.C., 2012. Krüppel-like factor 4 is involved in cell scattering induced by hepatocyte growth factor. J. Cell Sci. 125, 4853–4864. https://doi.org/10.1242/jcs.108910

Leka-Emiri, S., Chrousos, G.P., Kanaka-Gantenbein, C., 2017. The mystery of puberty initiation: genetics and epigenetics of idiopathic central precocious puberty (ICPP). J. Endocrinol. Invest. https://doi.org/10.1007/s40618-017-0627-9

Mahoney, J.E., Mori, M., Szymaniak, A.D., Varelas, X., Cardoso, W. V., 2014. The Hippo Pathway Effector Yap Controls Patterning and Differentiation of Airway Epithelial Progenitors. Dev. Cell 30, 137–150. https://doi.org/10.1016/j.devcel.2014.06.003

Mahony, S., Benos, P. V, 2007. STAMP: a web tool for exploring DNA-binding motif similarities. Nucleic Acids Res. 35, W253–8. https://doi.org/10.1093/nar/gkm272

Martin, L.J., Tremblay, J.J., 2009. The nuclear receptors NUR77 and SF1 play additive roles with c-JUN through distinct elements on the mouse Star promoter. J. Mol. Endocrinol. 42, 119–129. https://doi.org/10.1677/JME-08-0095

Matys, V., Fricke, E., Geffers, R., Gössling, E., Haubrock, M., Hehl, R., Hornischer, K., Karas, D., Kel, A.E., Kel-Margoulis, O. V, Kloos, D.-U., Land, S., Lewicki-Potapov, B., Michael, H., Münch, R., Reuter, I., Rotert, S., Saxel, H., Scheer, M., Thiele, S., Wingender, E., 2003. TRANSFAC: transcriptional regulation, from patterns to profiles. Nucleic Acids Res. 31, 374–8.

Meng, Z., Moroishi, T., Guan, K.L., 2016. Mechanisms of Hippo pathway regulation. Genes Dev. https://doi.org/10.1101/gad.274027.115

Mesrouze, Y., Aguilar, G., Bokhovchuk, F., Martin, T., Delaunay, C., Villard, F., Meyerhofer, M., Zimmermann, C., Fontana, P., Wille, R., Vorherr, T., Erdmann, D., Furet, P., Scheufler, C., Schmelzle, T., Affolter, M., Chène, P., 2020. A new perspective on the interaction between the Vg/VGLL1-3 proteins and the TEAD transcription factors. Sci. Rep. 10, 1–12. https://doi.org/10.1038/s41598-020-74584-x

Mizuno, T., Murakami, H., Fujii, M., Ishiguro, F., Tanaka, I., Kondo, Y., Akatsuka, S., Toyokuni, S., Yokoi, K., Osada, H., Sekido, Y., 2012. YAP induces malignant mesothelioma cell proliferation by upregulating transcription of cell cycle-promoting genes. Oncogene 31, 5117–5122. https://doi.org/10.1038/onc.2012.5

Mobley, K.B., Aykanat, T., Czorlich, Y., House, A., Kurko, J., Miettinen, A., Moustakas-Verho, J., Salgado, A., Sinclair-Waters, M., Verta, J.P., Primmer, C.R., 2021. Maturation in Atlantic salmon (Salmo salar, Salmonidae): a synthesis of ecological, genetic, and molecular processes. Rev. Fish Biol. Fish. https://doi.org/10.1007/s11160-021-09656-w

Muñoz-Cueto, J.A., Zmora, N., Paullada-Salmerón, J.A., Marvel, M., Mañanos, E., Zohar, Y., 2020. The gonadotropin-releasing hormones: Lessons from fish. Gen. Comp. Endocrinol. https://doi.org/10.1016/j.ygcen.2020.113422

Obayashi, T., Kagaya, Y., Aoki, Y., Tadaka, S., Kinoshita, K., 2019. COXPRESdb v7: a gene coexpression database for 11 animal species supported by 23 coexpression platforms for technical evaluation and evolutionary inference. Nucleic Acids Res. 47, D55–D62. https://doi.org/10.1093/nar/gky1155

Oku, Y., Nishiya, N., Tazawa, T., Kobayashi, T., Umezawa, N., Sugawara, Y., Uehara, Y., 2018. Augmentation of the therapeutic efficacy of <scp>WEE</scp> 1 kinase inhibitor <scp>AZD</scp> 1775 by inhibiting the <scp>YAP</scp> –E2F1–<scp>DNA</scp> damage response pathway axis. FEBS Open Bio 8, 1001–1012. https://doi.org/10.1002/2211-5463.12440

Pacini, V., Petit, F., Querat, B., Laverriere, J.N., Cohen-Tannoudji, J., L’Hôte, D., 2019. Identification of a pituitary ERα-activated enhancer triggering the expression of Nr5a1, the earliest gonadotrope lineage-specific transcription factor. Epigenetics and Chromatin 12, 1–17. https://doi.org/10.1186/s13072-019-0291-8

Pfaffl, M.W., 2001. A new mathematical model for relative quantification in real-time RT-PCR. Nucleic Acids Res. 29, e45.

Pohl, J., Golovko, O., Carlsson, G., Örn, S., Schmitz, M., Ahi, E.P., 2021. Gene co-expression network analysis reveals mechanisms underlying ozone-induced carbamazepine toxicity in zebrafish (Danio rerio) embryos. Chemosphere 276, 130282. https://doi.org/10.1016/j.chemosphere.2021.130282

Riddick, G., Kotliarova, S., Rodriguez, V., Kim, H.S., Linkous, A., Storaska, A.J., Ahn, S., Walling, J., Belova, G., Fine, H.A., 2017. A Core Regulatory Circuit in Glioblastoma Stem Cells Links MAPK Activation to a Transcriptional Program of Neural Stem Cell Identity. Sci. Rep. 7, 1–15. https://doi.org/10.1038/srep43605

Riverso, M., Montagnani, V., Stecca, B., 2017. KLF4 is regulated by RAS/RAF/MEK/ERK signaling through E2F1 and promotes melanoma cell growth. Oncogene 36, 3322–3333. https://doi.org/10.1038/onc.2016.481

Sinclair-Waters, M., Piavchenko, N., Ruokolainen, A., Aykanat, T., Erkinaro, J., Primmer, C.R., 2021. Refining the genomic location of single nucleotide polymorphism variation affecting Atlantic salmon maturation timing at a key large-effect locus. Mol. Ecol. https://doi.org/10.1111/mec.16256

Szklarczyk, D., Franceschini, A., Wyder, S., Forslund, K., Heller, D., Huerta-Cepas, J., Simonovic, M., Roth, A., Santos, A., Tsafou, K.P., Kuhn, M., Bork, P., Jensen, L.J., von Mering, C., 2015. STRING v10: protein-protein interaction networks, integrated over the tree of life. Nucleic Acids Res. 43, D447–52. https://doi.org/10.1093/nar/gku1003

Thissen, D., Steinberg, L., Kuang, D., Thissen, D., Steinberg, L., Kuang, D., 2002. Quick and Easy Implementation of the Benjamini-Hochberg Procedure for Controlling the False Positive Rate in Multiple Comparisons. J. Educ. Behav. Stat. 27, 77–83.

Varlinskaya, E.I., Vetter-O’Hagen, C.S., Spear, L.P., 2013. Puberty and gonadal hormones: Role in adolescent-typical behavioral alterations. Horm. Behav. https://doi.org/10.1016/j.yhbeh.2012.11.012

Verta, J.P., Debes, P.V., Piavchenko, N., Ruokolainen, A., Ovaskainen, O., MoustakasVerho, J.E., Tillanen, S., Parre, N., Aykanat, T., Erkinaro, J., Primmer, C.R., 2020. Cis-regulatory differences in isoform expression associate with life history strategy variation in Atlantic salmon. PLoS Genet. 16, e1009055. https://doi.org/10.1371/journal.pgen.1009055

Whitlock, K.E., Postlethwait, J., Ewer, J., 2019. Neuroendocrinology of reproduction: Is gonadotropin-releasing hormone (GnRH) dispensable? Front. Neuroendocrinol. https://doi.org/10.1016/j.yfrne.2019.02.002

Whitlock, N.C., Bahn, J.H., Lee, S.H., Eling, T.E., Baek, S.J., 2011. Resveratrol-induced apoptosis is mediated by early growth response-1, Krüppel-like factor 4, and activating transcription factor 3. Cancer Prev. Res. 4, 116–127. https://doi.org/10.1158/1940-6207.CAPR-10-0218

Wu, Y., Ma, S., Xia, Y., Lu, Y., Xiao, S., Cao, Y., Zhuang, S., Tan, X., Fu, Q., Xie, L., Li, Z., Yuan, Z., 2017. Loss of GCN5 leads to increased neuronal apoptosis by upregulating E2F1- and Egr-1-dependent BH3-only protein Bim. Cell Death Dis. 8, e2570–e2570. https://doi.org/10.1038/cddis.2016.465

Zhang, Y., Xu, N., Xu, J., Kong, B., Copple, B., Guo, G.L., Wang, L., 2014. E2F1 is a novel fibrogenic gene that regulates cholestatic liver fibrosis through the Egr-1/SHP/EID1 network. Hepatology 60, 919–930. https://doi.org/10.1002/hep.27121

Zheng, C., Ren, Z., Wang, H., Zhang, W., Kalvakolanu, D. V., Tian, Z., Xiao, W., 2009. E2F1 induces tumor cell survival via nuclear factor-κB-dependent induction of EGR1 transcription in prostate cancer cells. Cancer Res. 69, 2324–2331. https://doi.org/10.1158/0008-5472.CAN-08-4113

Zhu, C., Li, L., Zhang, Z., Bi, M., Wang, H., Su, W., Hernandez, K., Liu, P., Chen, J., Chen, M., Huang, T.H.M., Chen, L., Liu, Z., 2019. A Non-canonical Role of YAP/TEAD Is Required for Activation of Estrogen-Regulated Enhancers in Breast Cancer. Mol. Cell 75, 791-806.e8. https://doi.org/10.1016/j.molcel.2019.06.010

